# Modeling synthetic serum marker kinetics for monitoring deep-tissue gene expression

**DOI:** 10.1101/2025.11.17.688787

**Authors:** Nicolas Buitrago, Josefina Brau, Jerzy O. Szablowski

**Author notes:** Correspondence should be addressed to J.O.S.

## Abstract

Serum markers could theoretically enable monitoring of gene expression dynamics with a simple blood draw. However, such markers are typically used to measure long-term changes, such as the progression of disease over multiple days or months. In this theoretical study, we determine the maximum theoretical temporal resolution and precision – the ability to distinguish rapid changes in gene expression and to obtain the true frequency of such changes, respectively. As a model of such processes, we used Released Markers of Activity (RMAs) - a class of synthetic serum markers that are expressed in neurons in the brain but transported into the blood. RMAs are orthogonal to physiological processes and have tunable levels, production rate, and onset time, providing well-defined data for serum marker production, tissue transport, and detection. We explore several scenarios where RMAs were used, including monitoring tissue transduction, changes in endogenous gene expression, and drug-induced marker expression. We demonstrate that the temporal resolution of monitoring primarily depends on the extrinsic noise level of the RMA signal and protein serum half-life. Additionally, we find that decreasing the serum half-life results in improved temporal precision at the cost of signal intensity. To enable broad use of this model, we developed a library and interface for running simulations to approximate marker trajectories to inform experimental design decisions and optimize marker detection.

**Author Summary:** Gene expression is a foundational driver of biological processes. We use synthetic serum markers that can report on gene expression with a blood draw. These synthetic markers have well-defined parameters and thus form a useful model for understanding the maximum theoretical temporal resolution and precision of monitoring gene expression. These two variables describe, respectively, the fastest changes in gene expression and how accurately the kinetics can be reflected through a serum marker measurement. Developing more robust reporter systems requires understanding how biological properties of these markers, such as serum half-life or measurement noise, affect their kinetics. Here, we developed computational models describing *in vivo* behavior of synthetic serum markers to identify the most critical parameters for optimization. We show that serum half-life and measurement noise are the main contributors to the temporal precision and resolution of monitoring. Additionally, we demonstrate the ability to predict marker levels in various contexts, including constitutive and drug-induced expression.

## Introduction

Monitoring levels of serum markers are among the most commonly used test in medicine. Thanks to such measurements, one can learn about the physiology of deep tissues with a simple blood draw. Protein-based markers are the result of gene expression, making them an attractive interface for tracking the dynamics of gene expression in deep tissues over time. However, serum markers are generally used to track long-term changes in tissue physiology, such as the development of disease [1–5].

Synthetic serum markers with defined production rates, half-lives, and mechanism of transport from the tissue into the blood are an attractive model system for studying the kinetics of gene expression underlying serum marker levels. In some cases, markers can diffuse into the circulation without involvement of an intracellular transport receptor using fenestrated endothelium, e.g., in the liver [6] or in the brain following temporary opening of the blood-brain barrier (BBB) [7]. A more generalizable case involves markers undergoing receptor-mediated reverse transcytosis from the tissue into the blood, with a prime example being an intact BBB for antibodies that exit the brain [8]. Recently, synthetic serum markers called released markers of activity (**RMAs**) were engineered to track endogenous and drug-induced gene expression in the brain, enabling real-time and repeated *in vivo* monitoring [9]. RMAs are fully genetically encodable reporters with a Gaussia luciferase (Gluc) and fragment crystallizable (Fc) domain of immunoglobulin G (IgG) antibodies, enabling their transport across the BBB into circulation [9–12]. The Fc region has also been shown to contribute to a prolonged serum half-life [9,13], which may affect the temporal resolution or precision for monitoring rapidly changing gene expression. The fast erasable RMA variant also exists and improves the off kinetics by enabling on-demand clearance from the serum. By introducing a proteolytic cleavage site between the Gluc and Fc regions [14], the luciferase signal can be rapidly cleared by administration of a sequence-specific protease. However, an open question remains regarding how the production rates, marker levels, half-life, and other parameters affect the theoretical temporal resolution of RMAs and serum markers in general.

Here, we developed computational models to explore parameters relevant for optimizing reporter detection and temporal resolution. We specifically investigated the effects of reverse transcytosis, overall production, and degradation of RMAs in various contexts, including constitutive and drug-induced expression. We also evaluated the theoretical temporal resolution and precision of monitoring rapidly changing gene expression with RMAs. We define these variables as the ability to distinguish rapid changes in gene expression and the ability to accurately match the frequency of the RMA signal in the blood to the frequency of the target gene expression in the tissue of interest. Overall, we show that simulated RMA trajectories accurately reproduce experimental observations and that protein serum half-life is among the most significant properties affecting temporal resolution and precision. Additionally, we developed a graphical interface for simulating RMA trajectories under various conditions to better inform experimental design decisions, such as promoter strengths, sampling frequencies, and optimal time windows for monitoring gene expression with RMAs. These models provide a framework for predicting and interpreting RMA measurements in diverse gene expression monitoring applications, thereby improving our understanding of the temporal patterns of gene expression that can be feasibly monitored with serum markers in general.

## Results

We first developed a deterministic model of RMA expression driven by a constitutive promoter, such as the neuron-specific human synapsin 1 (hSyn). Here, we consolidate transcription, translation, and protein secretion into a single production term (**Fig 1a**). In this model, we make three core assumptions: 1) Transcription and translation kinetics are not rate limiting at the relevant time scales for RMA monitoring (days to weeks) [7,9,14,15]; 2) Dilution of the RMA protein by cell division is negligible in neuronal cell types, which are largely post-mitotic [16,17]; and 3), RMA efflux from brain to blood occurs primarily by receptor-mediated reverse transcytosis [10,18,19]. We fit this model to empirical RMA levels measured at 0-, 2-, and 3-weeks post-delivery of viral vectors harboring the RMA coding sequence (AAV-hSyn-Gluc-RMA-IRES-GFP) to either the CA1 region of the hippocampus, caudate putamen, or substantia nigra [9] (**Fig 1b**). RMA reverse transcytosis and degradation rates were constrained based on values reported for IgG antibody reverse transcytosis rates [10,18,20] and RMA serum half-life measurements [9]. Estimated values for RMA production, reverse transcytosis, and degradation were 1.38 × 10^-1^ ± 5.67 × 10^-2^ nM hr^-1^, 6.48 × 10^-1^ ± 7.75 × 10^-2^ hr^-1^, and 7.24 × 10^-3^ ± 2.23 × 10^-3^ hr^-1^ (mean ± std.), respectively (**Table S1**). To determine the relative importance of RMA production (𝑘_𝑅𝑀𝐴_), reverse transcytosis (𝑘_𝑅𝑇_), and degradation (𝛾_𝑅𝑀𝐴_) rates, we performed a global sensitivity analysis on the serum RMA signal using the method of Morris [21–23] with parameter ranges ± 50% from the estimated nominal values. The normalized absolute mean, µ*_norm_, and standard deviation, σ_norm_, of the elementary effects were used to rank the relative importance and nonlinearity or interactions of each parameter, respectively. We observed comparable relative importance and non-additive effects between production (µ*_norm_= 1.00 ± 0.043, σ_norm_ = 1.0) and degradation (µ*_norm_ = 0.73 ± 0.031, σ_norm_ = 0.85) rates at the 3-week timepoint (**Fig 1c-d**, µ*_norm_ = normalized absolute mean ± 95% confidence interval, σ_norm_ = normalized std.). Changes in reverse transcytosis rates within expected values for IgG antibodies were negligible (µ*_norm_ = 9.88 × 10^-4^ ± 1.10 × 10^-4^, σ = 2.65 × 10^-3^) for RMA proteins at 3 weeks post viral vector delivery, due to the much faster timescale compared to protein production and degradation. However, the reverse transcytosis rate was more important than protein degradation earlier on in the expression time course (**Fig S1**). Therefore, the reverse transcytosis parameter was not eliminated from this model to allow for generalizability to other RMA variants or serum markers with much shorter half-lives on the scale of minutes to hours, such as fast-erasable RMAs [14].

**Fig 1.**
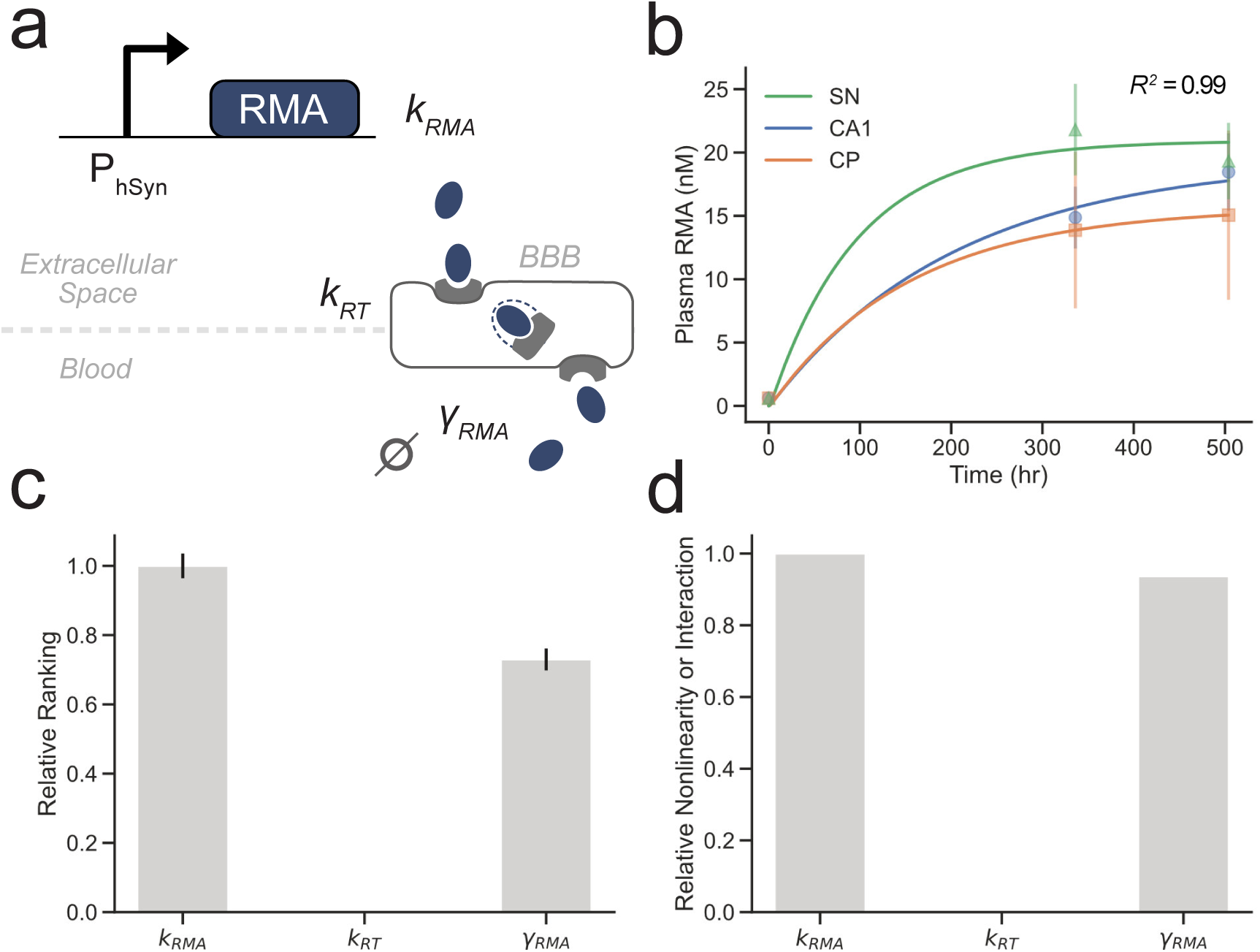
Constitutive RMA Kinetics. **a)** Model schematic for RMA production driven by a constitutive human synapsin promoter, receptor-mediated transport to the blood, and RMA degradation. **b)** Simulated RMA trajectories (solid lines) fit to experimental data (points, mean ± standard deviation) at 0, 2, and 3 weeks after delivery of AAV-hSyn-Gluc-RMA-IRES-GFP to the CA1 region of the hippocampus, caudate putamen (CP), or substantia nigra (SN) (average R^2^ = 0.98). **c)** Relative parameter importance and **d**) nonlinear or interaction effects for production, reverse-transcytosis, and degradation rates with respect to serum RMA.

To further investigate the temporal resolution and precision of RMA dynamics, we modified the constitutive RMA model to use an oscillating production term to test the monitoring of rapid changes in target gene expression (**Fig 2a**). We simulated RMA trajectories with production rates oscillating at a frequency of 1/72 hr^-1^ and varying half-life between 12.5 and 100 hours (**Fig 2b**). Both protein half-life and production frequency showed negative correlation with dynamic range (**Fig 2c**) [24]. Thus, while increasing RMA serum half-life allows for reporter accumulation and greater maximum serum concentration, it negatively alters the dynamic range of measurement (**Fig 2d-e**). To explore the limits of temporal resolution with respect to half-life, production oscillation frequency, and extrinsic noise levels, we introduced artificial noise in the plasma signal. We added homoscedastic Gaussian noise with varying standard deviations between 0.01 and 0.2 to the solution of the deterministic model, simulating measurement noise, and computed the temporal precision and resolution (**Fig 2f**). A total of 1000 trajectories were generated with added random noise, and the fraction of recovery of the matching input frequency was calculated for varying frequencies from 1/30 to 1/3 hr^-1^. To determine whether the correct input frequency was recovered from the RMA signal, we computed the power spectral density (PSD) of the noisy RMA signal using the Welch method and estimated the fundamental frequency from the peak of the PSD. We defined a true recovery of the input frequency as a fundamental frequency within 5% error and a signal-to-noise ratio of 2:1. We repeated this process for various degradation rates and found greater recovery of higher frequencies with decreasing serum half-life (**Fig 2g**) at the cost of reduced signal intensity. To investigate the impulse response of RMA dynamics with varying noise levels, we compute the power at the fundamental frequency as a fraction of the total power of the noisy RMA signal. We found that the fractional power at the input gene expression frequency decreases rapidly with increasing standard deviation of the added Gaussian noise (**Fig 2h**). Additionally, we examined the coherence [25], or relationship between the input gene expression oscillation and the serum RMA signal, with respect to the simulated measurement noise level. We observed a decrease in coherence with increased noise, as expected, and found that lower oscillation frequencies were more tolerant to higher noise levels (**Fig 2i**). Finally, we performed a global sensitivity analysis on the temporal precision, temporal resolution, and coherence with respect to RMA production, reverse transcytosis, degradation, and noise levels. We show that while the RMA degradation rate is important for recovering precise frequencies of input gene expression activity (**Fig 2j**), the temporal resolution and coherence primarily depend on the noise levels of the system (**Fig 2k-l**).

**Fig 2.**
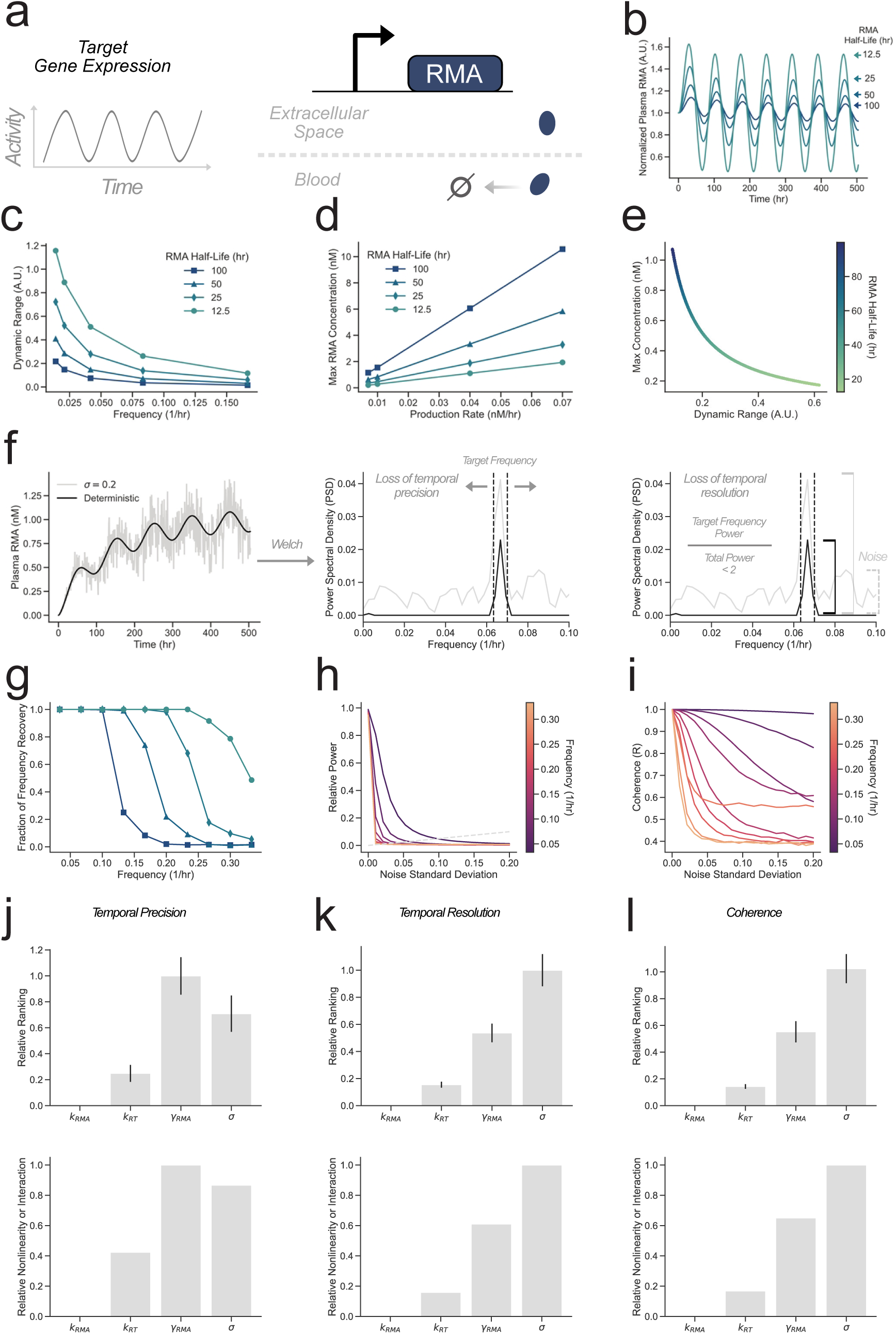
Temporal Precision and Resolution of RMA Dynamics. **a)** An arbitrary oscillating target gene controls the activity of RMA production. **b)** Normalized serum RMA signal with varying half-lives from 12.5 to 100 hours and an oscillating production term with frequency 1/72 hr^-1^. **c)** Dynamic range of normalized plasma RMA signal with respect to production frequency. **d)** Theoretical maximum RMA concentration with varying production rate and serum half-life. **e)** Relationship between maximum RMA concentration and dynamic range with varying serum half-life. **f**). Oscillating RMA dynamics with added Gaussian noise with standard deviation of 0.2 over a 3-week time course. The power spectral density of RMA signals is used to estimate the temporal precision and resolution given the frequency of the target gene driving the expression of RMAs. **g)** The fraction of recovery of the fundamental frequency from noisy RMA signals with added Gaussian noise (σ = 0.05) over a 3-week time course. **h)** Relative power at the fundamental frequency used in the production term and **i)** coherence with respect to noise strength. Relative parameter importance and nonlinear or interaction effects for production rate, reverse transcytosis rate, degradation rate, and noise strength with respect to **j)** temporal precision, **k)** temporal resolution, **l)** and coherence.

Inducible gene expression may be used in various contexts where a reporter protein is produced in response to specific stimuli. To further evaluate RMAs in this context, we used a simple inducible expression system where RMA production is driven by a constitutively expressed tetracycline transcriptional activator (tTA) in the absence of tetracycline or tetracycline derivatives such as doxycycline (**Fig 3a**). We specifically focused on the case where removal of doxycycline induces RMA expression, a commonly used scenario in neuroscience [26,27]. Thus, we assume that doxycycline concentrations reach a steady state before the beginning of the simulation and suppress RMA expression. Additionally, we include leaky expression of RMAs even in the presence of doxycycline. We simulated RMA trajectories with increasing doxycycline concentrations up to 625 mg/kg food [28] and assumed 4.5 g/day food intake for a 30g C56BL/6J mouse (**Fig 3b**) [29]. A global sensitivity analysis was repeated for the Tet-inducible model to evaluate the relative importance of RMA, doxycycline, and tTA parameters on the serum RMA levels. In many experiments, RMA levels are measured at least 24 hours after withdrawing doxycycline [9,14,15]. Thus, we began by examining the importance of each parameter on serum RMA levels at 24 hours post-doxycycline withdrawal. We found that RMA serum levels were most sensitive to the tTA-TetO binding dissociation constant and RMA serum degradation rate early on during doxycycline clearance (**Fig 3c,e**). We also evaluated the sensitivity of serum RMA levels with respect to each parameter at a second commonly used timepoint of 3 weeks post doxycycline withdrawal [9,14,15] (**Fig 3d,f**), finding a similar dependence of RMA serum levels on the tTA-TetO binding dissociation constant. However, that parameter was followed by the RMA degradation rate as well.

**Fig 3.**
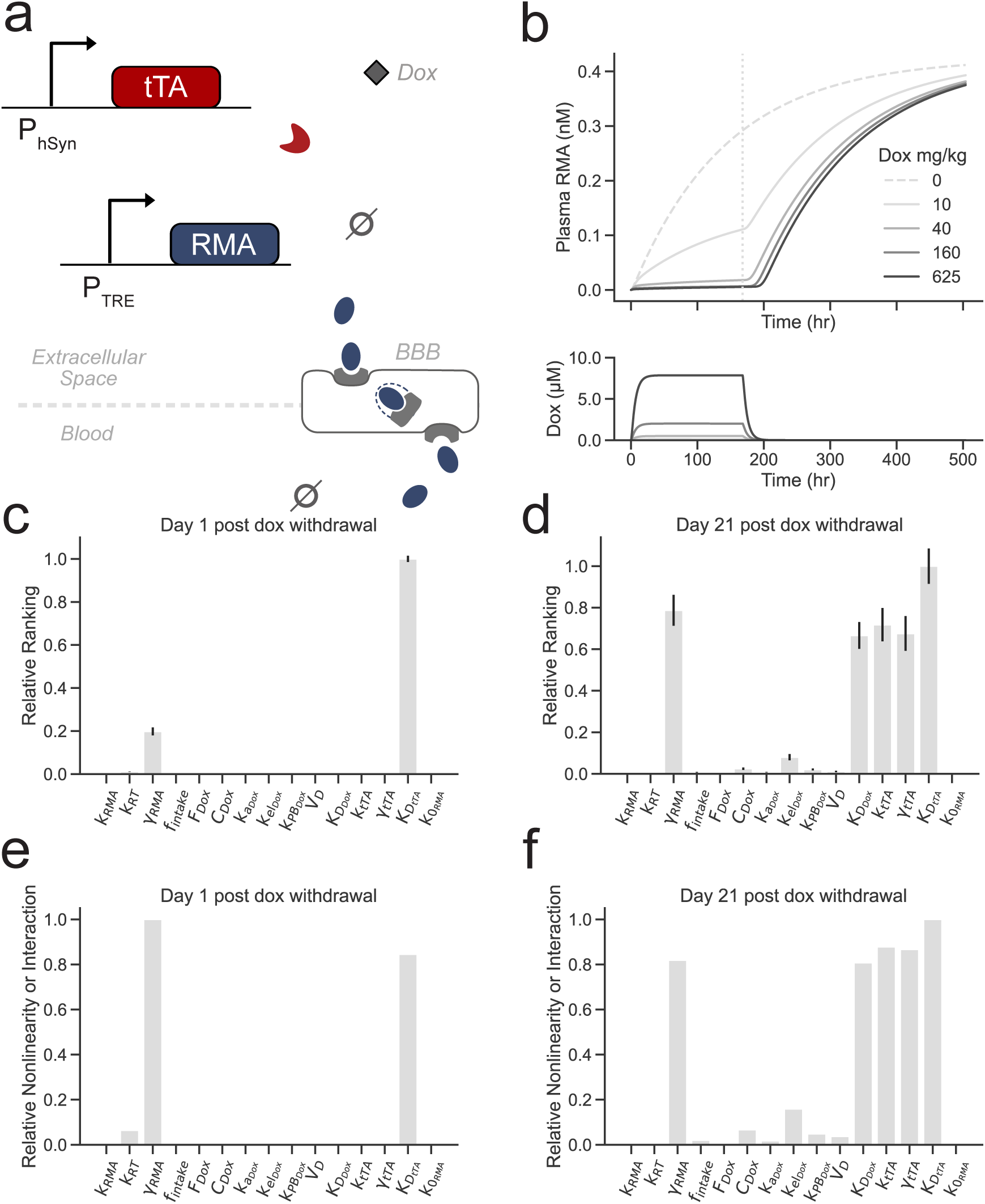
Inducible RMA kinetics. **a)** Model scheme for RMA production induced by a constitutively expressed tTA in the absence of doxycycline. **b)** Simulated RMA trajectories with doxycycline withdrawal on day 7 (vertical dashed line) at varying doxycycline dose from 0 to 625mg/kg food and average daily food intake of 4.5g for a 30g C57BL/6J mouse. **c)** Relative ranking of parameter importance on day 1 or **d)** day 21 after doxycycline withdrawal and **e-f)** nonlinear or or interaction effects with respect to serum RMA levels.

To better evaluate the effects of neuronal activity on RMA kinetics, we developed a model based on the synthetic gene circuit described in Lee et al. 2024 [9]. Here, an excitatory synthetic receptor hM3Dq is constitutively expressed in the brain under a human synapsin promoter [30–32]. Administration of hM3Dq’s corresponding ligand, such as clozapine-N-oxide (CNO), results in conversion to its parent compound, clozapine (CLZ), and Gq signaling leading to increased neuronal activity [33,34]. Subsequently, the synthetic robust activity marking (RAM) promoter is induced by neural activity [35] and drives the production of tTA, which then drives the expression of RMA. Thus, tTA and RMA may be expressed in response to DREADD-induced neural activity in the absence of doxycycline (**Fig 4a**). We fitted our model to RMA levels measured at 0, 24, and 48 hours after intraperitoneal injection of 1mg/kg CNO (**Fig 4b**). Kinetic parameters for CNO and CLZ dynamics were fitted to experimental measurements and fixed in the complete chemogenetic model (**Fig S2**) [33]. We then simulated plasma RMA, brain CNO, CLZ, and tTA levels after administration of 2.5mg/kg CNO using the same parameters estimated from the lower dose group (average *R^2^* = 0.89, **Table S4**). Peak brain tTA concentration was 38.56 and 42.81 nM at 1mg/kg and 2.5mg/kg CNO, respectively. The peak CNO and CLZ brain concentrations were 1.86 and 148.41 nM at 1mg/kg CNO and 4.66 and 371.06 nM at 2.5mg/kg CNO. The calculated plasma RMA levels reached 18.62 and 18.71 nM for 1 and 2.5mg/kg CNO, respectively, at 48 hours post-intraperitoneal injection. Finally, we fixed doxycycline and CNO/CLZ kinetic parameters, including absorption and clearance rates, and performed a global sensitivity analysis on the remaining free parameters listed in **Table S4** (**Fig 4c-d**). We found that RMA production showed the highest relative importance on serum RMA levels in this model, followed by the tTA-TetO binding dissociation constant at the 48-hour time point, consistent with the constitutive expression case where RMA production is the main driver of variability at earlier time points. Interestingly, hM3Dq steady state concentration and CNO or CLZ EC_50_ were among the lowest-ranked parameters screened in this system.

**Fig 4.**
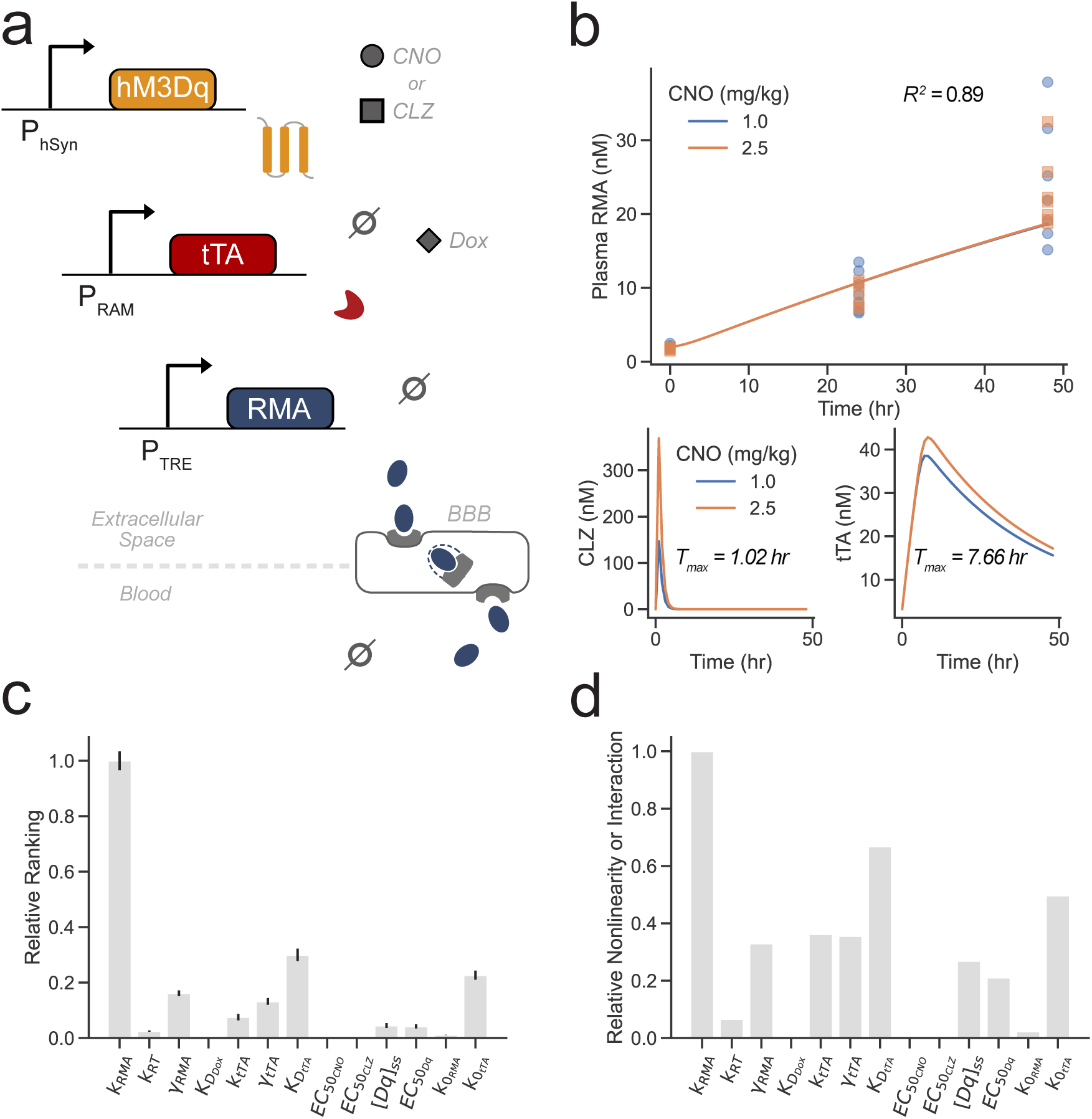
Chemogenetic Activated RMA with Tet-Off Gating. **a)** Model scheme of constitutively expressed hM3Dq driving the production of tTA upon CNO administration. Free tTA unbound to doxycycline drives the expression of RMA. **b)** RMA serum levels (solid lines) fit to experimental measurements (points) after intraperitoneal injection of either 1 (blue) or 2.5mg/kg (orange) CNO. **c)** Relative parameter importance and **d)** nonlinear or interaction effects with respect to serum RMA level.

## Discussion

We developed three main models for predicting constitutive and drug-induced RMA expression. In all cases, we assumed that RMA transport into the circulating blood occurs mainly through receptor-mediated reverse transcytosis [10,18]. While other transport mechanisms were not considered, the constitutive and drug-induced models were consistent with empirical measurements [9]. Importantly, we note that in this work, measured relative light units (RLUs) from previous studies were converted to molar concentrations using their associated luciferase standard curve. However, there may be differences in RLU signal due to variation in luciferase substrate preparation, instrumentation, and sample type. Thus, converting RLUs from other experiments with the standard curve used here may not produce accurate concentrations. Further, we explored how properties such as serum half-life and overall noise of the system affect the temporal precision and resolution of monitoring. We demonstrate that decreasing the serum half-life can lead to improved temporal precision at the cost of lower absolute fold changes in serum RMA levels. Additionally, we show that temporal resolution mainly depends on the relative noise within the RMA signal.

For more complex systems that involve doxycycline inhibition, we found that the tTA-TetO binding dissociation constant, along with tTA production and degradation rates, had the greatest impact on model output. Thus, RMA dynamics may also be tuned using tTA variants such as d2tTA or engineering other tTAs for specific contexts and use with RMAs. This becomes increasingly important for RMA systems where tTA is driven by an RNA-based sensor targeting endogenous genes *in vivo* [15]. In the future, these models may be adapted further for predicting the dynamics of markers recovered by insonation [7] or RMAs driven by different synthetic promoters. Overall, these models can be further adapted and generalized to other serum markers expressed in peripheral tissues.

## Methods

### RMA Kinetic Models

For constitutive RMA expression, the production of RMA in the brain tissue and transport into the blood is described by a system of coupled ordinary differential equations

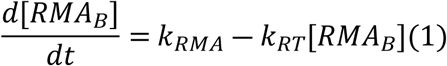

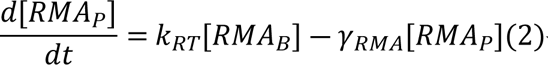

Where 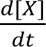 is the time derivative of the concentration for species 𝑋; [𝑅𝑀𝐴_𝐵_] and [𝑅𝑀𝐴_𝑃_] are the concentrations of RMA in the brain tissue and plasma, respectively; 𝑘_𝑅𝑀𝐴_, 𝑘_𝑅𝑇_, and 𝛾_𝑅𝑀𝐴_ are the RMA production, reverse transcytosis, and degradation rates, respectively.

As a proxy for monitoring rapidly changing gene expression with RMAs, we modify the constant production 𝑘_𝑅𝑀𝐴_ to a sinusoid oscillating production term

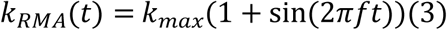

where 𝑘_𝑚𝑎𝑥_ is the max RMA production rate, 𝑓 is the ordinary frequency, and 𝑡 is time.

To expand to drug-inducible RMA expression with the Tet-Off system, we assume constitutive expression of the tetracycline transcriptional activator (tTA) and describe its dynamics by the differential equation

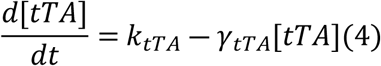

with tTA production and degradation rate constants 𝑘_𝑡𝑇𝐴_and 𝛾_𝑡𝑇𝐴_, respectively. The fraction of active tTA not bound to doxycycline (𝜃_𝑡𝑇𝐴_), is described by a Hill function,

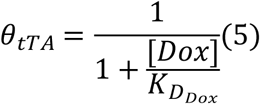

where 𝐾_𝐷𝐷𝑜𝑥_ is the equilibrium dissociation constant of tTA-Dox binding. A two-compartment model was used to describe the dox absorption, brain distribution, and elimination within a set administration period (**Table S2**). The production of RMA is then dependent on the concentration of free tTA and modified for leaky expression,

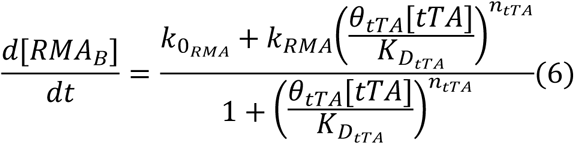

where 𝑘_0𝑅𝑀𝐴_ and 𝑘_𝑅𝑀𝐴_ are the leaky basal and max RMA production rates, respectively; 𝐾_𝐷𝑡𝑇𝐴_ and 𝑛_𝑡𝑇𝐴_ are the equilibrium dissociation constant of tTA-TetO operator binding and the tTA Hill coefficient, respectively. The remaining equations describing the change in brain and plasma RMA concentrations (**Eq. 1-2**) remain the same as in the constitutive model.

The final chemogenetic RMA model was based on the genetic circuit described in Lee et al. 2024 for monitoring drug-induced neural activity [9]. Here, we assume constitutive expression of the activator DREADD hM3Dq (referred to as Dq from here), which may be activated by clozapine (CLZ) or its metabolite clozapine-N-oxide (CNO) [30–33].

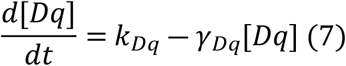

The dynamics of CNO and CLZ are described by a two-compartment model with first-order absorption (**Fig S2a**). The fraction of active hM3Dq at a given concentration of CNO or CLZ is then given as,

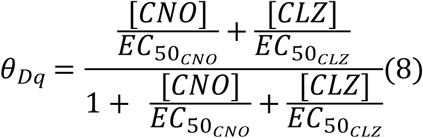

and the production of tTA is described by a Hill function modified for leaky production,

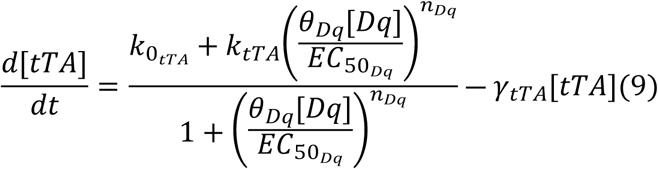

where 𝑘_0𝑡𝑇𝐴_ and 𝑘_𝑡𝑇𝐴_ are the leaky basal and max tTA production rates, respectively; 𝑛_𝐷𝑞_is the Dq activation Hill coefficient and 𝛾_𝑡𝑇𝐴_is the tTA degradation rate constant. The remaining equations describing the dynamics of doxycycline (**Eq. S2-4**) and RMAs (**Eq. 2, 6**) remain the same as the Tet-RMA model. All models were implemented in Python 3.13 and simulated using Jax 0.6.1 and Diffrax 0.7.0 for numerical approximations.

### Parameter Estimation

Models were fit to experimental measurements of plasma RMA luminescence converted to concentration using a luciferase standard curve [7]. Parameters were estimated by particle swarm optimization [36–38] using an objective function defined as,

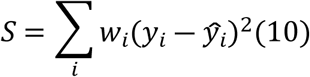

where 𝑤_𝑖_, 𝑦_𝑖_, and 𝑦_𝑖_are the weighting factor, observed, and predicted values of data point 𝑖, respectively. The initial conditions for the particle swarm were set based on reported values in the literature, if available [9,10,18]. For the Tet-RMA model, parameters for doxycycline dynamics were fixed based on values reported in the literature [39–42]. Similarly, CNO and CLZ kinetic parameters were estimated by fitting a two-compartment model to brain CNO and CLZ concentrations after intraperitoneal administration of CNO and later fixed in the full chemogenetic model (**Fig S2b-e**) [33,43]. The remaining parameters were estimated as described.

### Sensitivity Analysis

Global sensitivity analysis was performed for each model using the Morris method [21–23]. Briefly, 250 parameter trajectories were generated [21] and used to simulate serum RMA levels. The elementary effect for parameter x_k_ was calculated as the finite difference of the output with respect to a parameter step and finite step Δ, such that the parameter set 𝑋 = (𝑥_1_,𝑥_2_,…, 𝑥_𝑘_) + Δ is within the set parameter bounds.

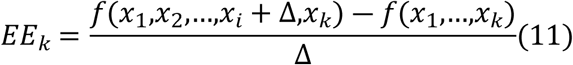

The absolute mean and standard deviation of the elementary effects for the k’th parameter for r replicates are defined as

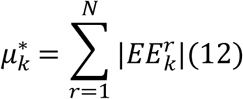

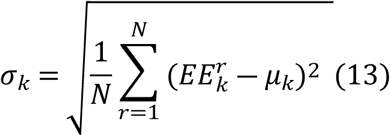

## Code Availability

All experimental data used in this study are published in the cited references. All source code and notebooks are available on GitHub at https://github.com/szablowskilab/rma-kinetics. Models are also available as a Python package on PyPI, and a web interface for running simulations is available at https://www.szablowskilab.org/rmamodel.

## Acknowledgements

This research was supported by the NIH Director’s New Innovator Award (DP2) to J.O.S. We thank Jonathon DeBonis and Dr. Oleg Igoshin for helpful discussions.

## Author Contributions

Conceptualization, validation, writing – original draft, review and editing: J.O.S., N.B.

Data curation: N.B., J.B.

Funding acquisition, Supervision: J.O.S.

Investigation, formal analysis, methodology, visualization, software, resources: N.B.

## Competing Interests

J.O.S. is a cofounder of Imprint Bio Inc and is a co-inventor on a patent describing the original RMA technology. The remaining authors declare no competing interest.

